# The strength of feedback processing is associated with resistance to visual backward masking during illusory contour processing in adult humans

**DOI:** 10.1101/2022.04.04.487043

**Authors:** John J. Foxe, Emily J. Knight, Evan J. Myers, Cody Zhewei Cao, Sophie Molholm, Edward G. Freedman

## Abstract

Re-entrant feedback processing is a key mechanism of visual object-recognition, especially under compromised viewing conditions where only sparse information is available and object features must be interpolated. Illusory contour stimuli are commonly used in conjunction with visual evoked potentials (VEP) to study these filling-in processes, with characteristic modulation of the VEP in the ~100-150ms timeframe associated with this re-entrant processing. Substantial inter-individual variability in timing and amplitude of feedback-related VEP modulation is observed, raising the question whether this variability might underlie inter-individual differences in the ability to form strong perceptual gestalts. Backward masking paradigms have been used to study inter-individual variance in the ability to form robust object perceptions before processing of the mask interferes with object-recognition. Some individuals recognize objects when the time between target object and mask is extremely short, whereas others struggle to do so even at longer target-to-mask intervals. We asked whether timing and amplitude of feedback-related VEP modulations were associated with individual differences in resistance to backward masking. Participants (N=40) showed substantial performance variability in detecting illusory contours at intermediate target-to-mask intervals (67ms and 117ms), allowing us to use *kmeans* clustering to divide the population into four performance groups (poor, low-average, high-average, superior). There was a clear relationship between the amplitude (but not the timing) of feedback-related VEP modulation and illusory contour detection during backward masking. We conclude that individual differences in the strength of feedback processing in neurotypical humans lead to differences in the ability to quickly establish perceptual awareness of incomplete visual objects.

## INTRODUCTION

The visual system can construct object gestalts from what are often highly fragmented or incomplete inputs (1). Contour-integration processes are an essential component of this ability, and are readily assayed in humans using the visual-evoked potential (VEP) technique. Considerable work has shown that VEP modulations due to contour-integration are measurable within 40-50 ms of initial afference in V1 and that these processes are, in large part, driven by feedback inputs from higher-order ventral visual stream regions, specifically within the lateral occipital complex (LOC) (2–7). Illusory contour stimuli are commonly used to interrogate contour integration processes, since equivalent stimulus features can be spatially configured such that they do, or do not, induce the perception of a contour. The brain responses to these stimuli can then be compared to investigate the spatio-temporal dynamics of contour integration (4). Work in non-human primates (8, 9) and mice (10) strongly supports a feedback model of contour integration, and transcranial magnetic stimulation in humans has been used to infer that illusory contour processes in hierarchically early visual cortex (i.e. V1 and V2) occur at a delay relative to those seen in LOC (11, 12). Thus, it can be assumed that illusory contour modulations reflect an iterative (or resonant) feedback process between LOC and early visual cortex (e.g., V1) (5). Across a series of studies by our research group, we have noted substantial inter-individual variability in the timing and robustness of VEP-derived illusory contour modulations, but the functional significance of these feedback-related differences has not been explored to our knowledge. However, in related work in clinical populations, it has been noted that delays in the onset of these processes are observed in young adults with schizophrenia (13), raising the possibility that delays in feedback processing have functional implications that may be related to well-established visual sensory processing deficits in this population (14–19). Here, we set out to establish whether the strength and timing of visual feedback processing during an illusory contour integration task would relate to variability in task performance and reflect the robustness of visual sensory-perceptual functioning.

To this end, an excellent tool in the hands of vision researchers for determining timing effects on visual processing is the so-called backward-masking paradigm (20–22). In this paradigm, presentation of a stimulus to be acted upon (hereafter referred to as the “target”), is rapidly followed by presentation of a second “masking” stimulus (the “mask”) that is intended to interfere with processing of the target. When the interval between target and mask is sufficiently rapid, perceptual discrimination of relevant features of the target can be fully extinguished (23–25). There is substantial inter-individual variability in susceptibility to backward masking (26), suggesting that there is variability in either the speed or the robustness of visual processing. A reasonable proposition is that those who show least susceptibility to backward masking may display the fastest and most robust processing of the initial target, such that by the time that neural signals associated with the mask are coursing through the visual system, sufficient target processing has already occurred to support the final perception.

More specifically, in the case of contour-integration, one might expect that those who show the earliest or most robust feedback processing signatures would be those with most resilience to the mask. The notion that backward masking can interfere with feedback processing was beautifully illustrated in a study in non-human primates by Lamme and colleagues, who studied figure-ground detection under backward masking conditions (27). In a considerable body of work, this research group showed that figure-ground segregation processes emerge in recordings of V1 neurons at about 80-100 ms following stimulus delivery, which is substantially later than the initial afferent input to V1 (8, 28), highly reminiscent of the illusory-contour effects reported in human studies (2, 4). In turn, these contextual modulations were selectively attenuated by anesthesia (29) consistent with the notion that they arose from feedback inputs from higher-order visual regions (30). In their 2002 study (27), Lamme and colleagues recorded the activity of V1 neurons in awake behaving macaques who were tasked with indicating when they detected figures from background. They showed that at relevant target-to-mask intervals, suppression of the characteristic feedback figure-ground modulations was clearly related to failure to discriminate the figures.

In a highly relevant study in human observers, the same research group recorded VEP data during a figure-ground discrimination task where they presented target stimuli containing figures and those containing no-figures while participants were required to simply indicate whether there was a figure present or not (22). In a no-mask condition, performance was near perfect and a clear and characteristic series of VEP modulations was evident beginning at about 110-140ms and again between 200-300ms, which were taken to reflect re-entrant (feedback) processing, consistent with their prior work in non-human primates. It is notable that these modulations in response to figure-ground stimulus configurations are highly similar to those observed to illusory contour (Kanizsa (31)) figures. In a second condition, the same target stimuli were followed immediately (i.e. no delay) by a highly effective masking stimulus and performance dropped to chance for all participants. Examining the VEP records, it became clear that the initial sensory VEP response was unaffected by the mask, whereas the biphasic figure-induced modulations that began at 110ms were completely absent. The authors reasonably concluded that the masking stimuli had specifically disrupted feedback processing and that this was why their participants could no longer discriminate figure-ground targets.

Following the logic of these Lamme studies, here we asked if the timing and robustness of feedback illusory contour processing, as measured using high-density electrophysiological mapping of the VEP, would be associated with resistance to the effects that backward masking has on perceptual performance in human observers. In an initial psychophysical experiment, we assessed participants’ abilities to recognize the presence or absence of illusory contour targets at various target-to-mask intervals, thereby stratifying our participants into four performance groups: (1) below average, (2) low average, (3) high average, and (4) superior performers. In a separate electrophysiological experiment, we measured the neural response to the presence versus absence of illusory contours in the same participants, for unmasked stimuli. A clear relationship between the robustness of the early illusory contour-related VEP modulation and resistance to the effects of rapid backward masking emerged, whether considering performance as a continuous variable or by performance group. These findings suggest that individual participant variance in the strength of cortical feedback during basic visual processing has tangible consequences for everyday perceptual function.

## METHODS

### Participants

40 participants (age 18-33 years (mean ± SD = 21 ±3 years)) were recruited to this study, and included 19 females and 21 males. All had normal or corrected-to-normal vision, normal hearing, and no history of neurologic disorder. All experimental procedures were approved by the institutional review board of the University of Rochester Medical Center. Written informed consent was obtained from all participants, and they were modestly compensated for their time in the laboratory ($15/hour).

### Stimuli

Stimuli were delivered using Presentation^®^ software (Version 18.0, Neurobehavioral Systems, Inc., Berkeley, CA, www.neurobs.com). A schematic of the experimental paradigm is presented in Figure 1 and we have made the electrophysiologic paradigm code freely available for download with appropriate attribution (32). Individuals were presented with a Kanisza figure containing four Pac-Man-like inducers (31). Each inducer occupied one of four corners equidistant from a central fixation point. The presence of an illusory contour (IC) was defined as alignment of the cut-out of the four inducers such that it collectively produced the image of a square. Conversely, the non-contour (NC) configuration existed when at least one of the cut-outs of the inducers was rotated away from the fixation point and did not produce the illusion of a completed square. The retinal eccentricity of the entire IC or NC image was set at 3.5° of visual angle. The support ratio (the ratio of the portion of the perimeter occupied by the inducers themselves to the total perimeter of the induced square shape) was held constant at 0.54. That is, 54% of the observed shape was completed by a real line. The paradigm included randomly intermixed eccentric and central presentations of these stimuli; however, only trials where stimuli were presented in the center of the visual field were included in the current analysis.

**Figure 1.**
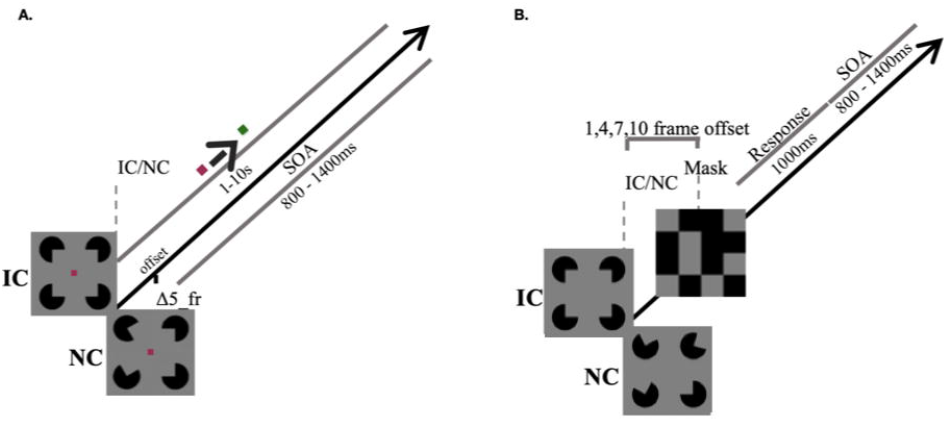
**A)** Schematic representation of the electrophysiologic experimental paradigm and timing. IC/NC stimuli appear on the screen for 80ms duration with 800-1400ms stimulus onset asynchrony (SOA). Central fixation point changes from red to green at variable intervals (1-10s), uncorrelated with IC/NC stimuli presentation. B) Schematic representation of the backward masking behavioral paradigm. IC or NC stimuli (50% probability of each configuration) were presented in the center of the screen for a duration of 1 frame (equivalent to ~17ms). Following the IC/NC offset, participants were presented with a gray screen with no stimuli for a randomized duration of either 1, 4, 7, or 10 frames prior to the presentation of a 100ms duration random black-gray checkerboard ‘mask’.

### Illusory Contour Electrophysiologic (EEG) Paradigm

For the EEG experiment, IC or NC stimuli were presented on the screen for a duration of 80ms with a jittered onset asynchrony between 800 – 1400ms. Participants engaged in an unrelated visual task, where they were instructed to focus on a central red fixation dot (4×4 pixels) against a gray background and to press a button on a game controller (SteelSeries 3GC USB 2.0) when the color changed to green (which occurred on average once every ten seconds and lasted approximately 160ms) on a random time course uncorrelated with IC/NC presentation (Figure 1A). The colors were chosen to be approximately isoluminant with the intention that the change in chromaticity was difficult to perceive without directly foveating. Participants were informed that additional objects would be presented on the monitor but instructed to “do your best to focus on the fixation dot.” There was no specific mention of the nature of the IC and NC stimuli included in the participant instructions. An eye-tracker (EyeLink 1000; SR Research) was used to ensure that participants were fixated on the center of the monitor. A 9-point calibration was performed prior to each test block which effectively paused stimulus onset when the participant’s eyes deviated > 5° from the center. Seven blocks, each containing approximately 60 stimuli with IC and NC stimuli intermixed and delivered in random order, were presented to each participant. Participants were allowed breaks as needed between blocks.

### Backward Masking Behavioral Task

For the behavioral experiment, the IC and NC stimuli, as well as equipment set-up, were identical to that described above. At a monitor refresh rate of 60Hz, IC or NC stimuli (50% probability of each configuration) were presented on the screen for a duration of 1 frame (equivalent to ~17ms). Following the IC/NC offset, participants were presented with a gray screen with no stimuli for a randomized duration of either 1, 4, 7, or 10 frames (approximately 17ms, 67ms, 117ms and 167ms respectively) prior to the presentation of a 100ms duration ‘mask’ (Figure 1B). The mask, generated using MATLAB® software (the MathWorks, Natick, MA, USA), consisted of a random black-gray checkerboard with each square equivalent in diameter to the IC/NC inducers. This mask was found during the piloting phase to be most effective at disrupting reported IC/NC perception as compared to other mask types employed in the literature (e.g. dots, line segments, and shapes (33)). There were additional trials consisting of mask alone or IC/NC not followed by the mask which served as controls. Participants were instructed to press a button on the game controller as fast as they could when they perceived the IC stimuli and to refrain from pressing any button when they did not perceive an IC.

### EEG Acquisition and Pre-processing

All participants sat in a sound-attenuated and electrically-shielded booth (Industrial Acoustics Company, The Bronx, NY) at a distance of 30 inches away from a monitor (Acer Predator Z35P 35” 21:9 100 Hz) 1280 x 1024-pixel resolution. Electroencephalographic (EEG) data were continuously recorded using a 64-channel Biosemi ActiveTwo acquisition system (Bio Semi B.V., Amsterdam, Netherlands). The setup included an analog- to-digital converter, and fiber-optic pass-through to a dedicated acquisition computer (digitized at 512 Hz; DC-to-150 Hz pass-band). EEG data were referenced to an active common mode sense electrode and a passive driven right leg electrode. EEG data were processed and analyzed offline using custom scripts that included functions from the EEGLAB (34) and ERPLAB Toolboxes (35) for MATLAB R2017a (the MathWorks, Natick, MA, USA). Raw data were filtered between 0.1 and 50 Hz using a Chebyshev type II filter and down-sampled to 250 Hz. Bad channels were automatically detected and interpolated using EEGLAB spherical interpolation. A summary table of the number of interpolated channels and rejected trials is shown in Table S1. Data were re-referenced to a frontal electrode (Fpz in the 10-20 system convention) and then divided into epochs that started 100ms before the presentation of each IC/NC stimulus and extended to 500ms post-stimulus onset. Trials containing severe movement artifacts or particularly noisy events were rejected if voltages exceeded ±50μV.

### Statistical Analyses

#### Behavioral Performance

Participant accuracy in detecting the presence of the IC stimulus during the Backward Masking psychophysics task was assessed according to d’= z(hit rate)-z(false alarm rate). Perfect hit rates of 1 and false alarm rates of 0 were corrected by −1/(2nIC) and +1/(2nNC), respectively, when present (36). Accuracy was determined for each of the five target-to-mask delay conditions (1, 4, 7, 10 frames and no mask). Per our main hypothesis, we observed significant individual variability in mask susceptibility and task performance, and developed a series of analyses to characterize these individual differences and their relationship to the electrophysiologic data. At the Δ1 frame delay, participants were generally unable to complete the task while by Δ10 frame delay all participants achieved d’ values >1. In contrast, at the Δ4 and Δ7 frame delays, individuals varied in their susceptibility to the mask. Because accuracy at the 4 and 7 frame conditions also conformed to a normal distribution, we then classified participants into four groups (below average, low average, high average, and superior) using a k-means clustering approach based on their performance at these two intermediate target-to-mask delays. Such classification allowed for sufficient averaging across participants within a group to visualize the relationship between VEP waveforms and task performance.

#### Electrophysiological data

To reduce the number of comparisons, analysis was limited to a pair of electrode sites over bilateral lateral occipital regions (PO3 and PO4 in the 10-20 system convention) where the IC-effect has been repeatedly shown to be maximal (37). Epochs were averaged to yield VEP to the IC and NC stimuli. A difference waveform representing the IC-effect was then calculated by subtracting the grand mean VEP to NC stimuli from the grand mean VEP to IC stimuli. The resulting distribution of activity showed a most pronounced IC-effect negative deflection between 100ms and 200ms, maximal at ~121ms on average, fully consistent with prior literature (4, 6). We assessed the within-participants difference in mean amplitude over a 10ms time window (116-126ms) for IC vs NC stimuli using Wilcoxon signed-rank tests with effect size determined by matched pairs rank biserial correlation (rc), due to violation of normality assumption.

To test the hypothesis that the speed and intensity of IC processing would be associated with the level of behavioral resistance to backward masking, we then defined a time window of 10ms centered around each individual participant’s IC-effect negative peak latency between 100ms-200ms to obtain a mean amplitude for the IC, NC, and IC-NC (IC-effect) waveforms for each individual. We conducted one-way analyses of variance (ANOVA) to assess the relationship of the four group performance classifications with IC-effect mean amplitude and IC-effect peak latency, followed by post-hoc Tukey procedure to evaluate significant interactions. Statistical analyses were performed in JASP (JASP Team (2019), Version 0.11.1). Finally, we assessed the relationship between electrophysiologic response and individual performance as a continuous factor using correlation analysis and linear regression to compare the d’ at each of the five target-to-mask delay conditions (1, 4, 7, 10 frames and no mask) and the IC-effect mean amplitude for each participant.

## RESULTS

### Backward Masking Task

Average accuracy in discrimination of the IC from NC stimuli, as measured by d’, for each target-to-mask delay condition, is represented in Figure 2A. Unsurprisingly, participants were unable to make accurate IC/NC discriminations beyond chance at Δ1 frame, the shortest target-to-mask interval. Accuracy was maximal in the no mask condition and decreased systematically with shorter target-to-mask delays from Δ10 frames to Δ1 frame. At the Δ10 frame delay, all participants demonstrated d’ measures >1. Consistent with the main hypothesis, the greatest inter-individual variability was observed at the Δ7 frame delay and the distribution of participant accuracy at the Δ4 and Δ7 target-to-mask delays conformed to a normal distribution (Figure 2B). As a result, participants were then grouped using a k-means clustering approach into four groups (below average, low average, high average, and superior) based on their performance at these two intermediate delays. There were no significant differences in age between the four performance-related groups. (Figure S1). Individual performance at each target-to-mask delay and group classification by accuracy (d’) at the Δ4 and Δ7 frames are depicted in Figure 2C-D. Individuals classified as below average, low average, high average, and superior performers had average accuracies of 0.53±0.25, 1.24±0.23, 1.76±0.36, and 2.90±0.53 at the Δ4 and 1.43±0.36, 2.57±0.52, 3.41±0.15, and 4.11±0.58, at the Δ7 frame delays, respectively.

**Figure 2.**
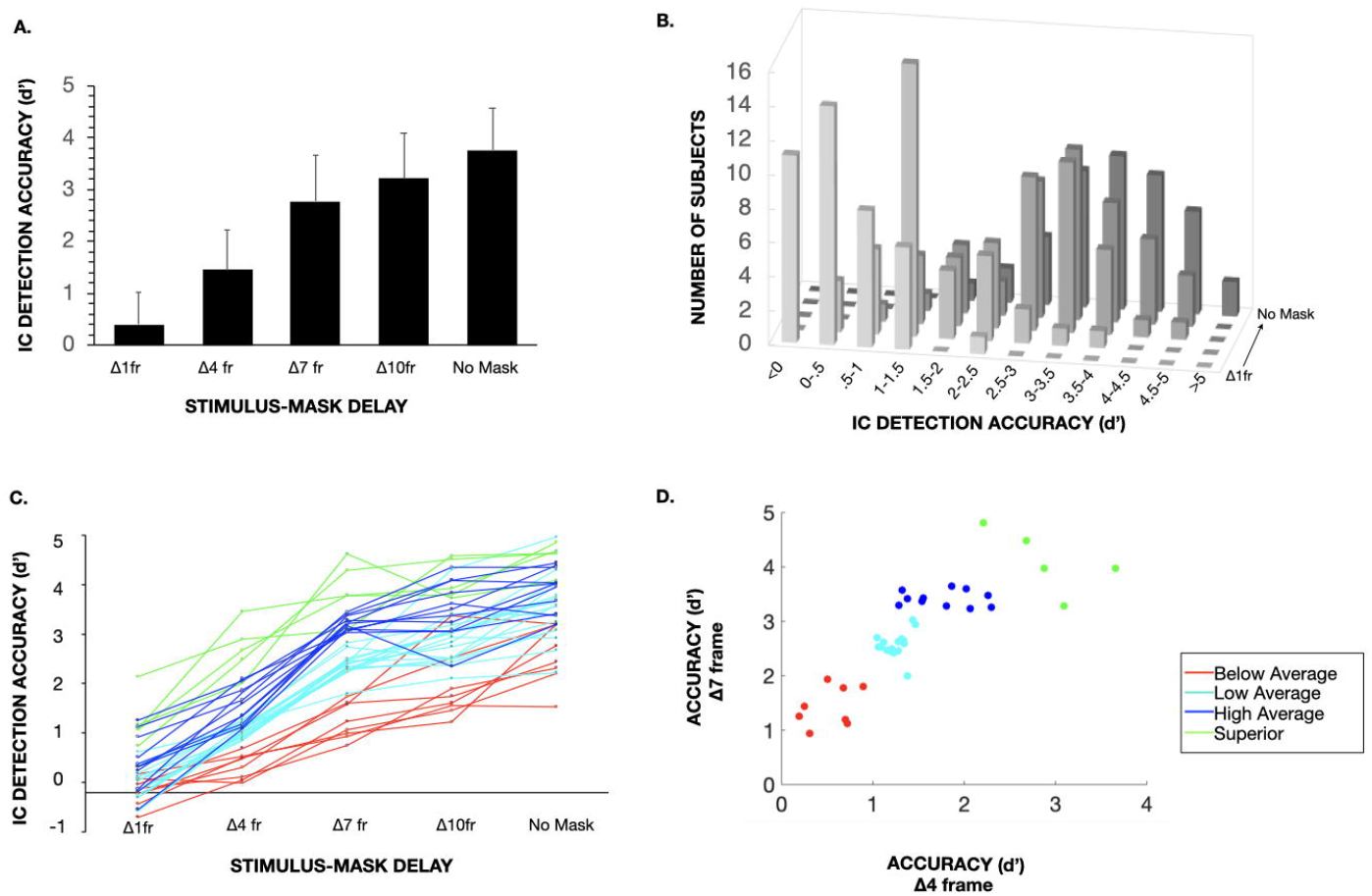
**A)** Whole group mean accuracy in IC detection as measured by d’ in each of the 1,4,7, and 10 frame stimulus-mask intervals and the “no mask” condition. Error bars = +1SD. B) Subjects binned by accuracy in IC detection as measured by d’ for each of the 1,4,7, and 10 frame stimulus-mask intervals and the “no mask” condition. **C)** Plot depicting increasing IC detection accuracy by stimulus-mask delay. Lines represent individual subjects with performance-related group classification indicated by color (red=below average, light blue=low average, dark blue=high average; green=superior). D) Cluster plot depicting individual subject accuracy at the 4 and 10 frame stimulus-mask delays with performance-related group classification as determined by *kmeans* clustering indicated by color (red = below average, light blue = low average, dark blue = high average; green = superior).

### Electrophysiologic Data

#### Grand Average IC-effect across All Participants

Figure 3 displays the grand-average VEP waveforms across all participants (irrespective of behavioral performance) elicited by the IC and NC stimuli, as well as the corresponding difference waves, over the left and right occipital regions (PO3 and PO4 in the 10-20 system convention). Consistent with previous literature, the evoked response to IC as compared to the NC stimuli was characterized by a more pronounced mean negativity over both the left (PO3: W=687, p=0.0002, rc=0.676) and right (PO4: W=669, p=0.0005, rc=0.632) occipital regions in the 116-126ms time window.

**Figure 3.**
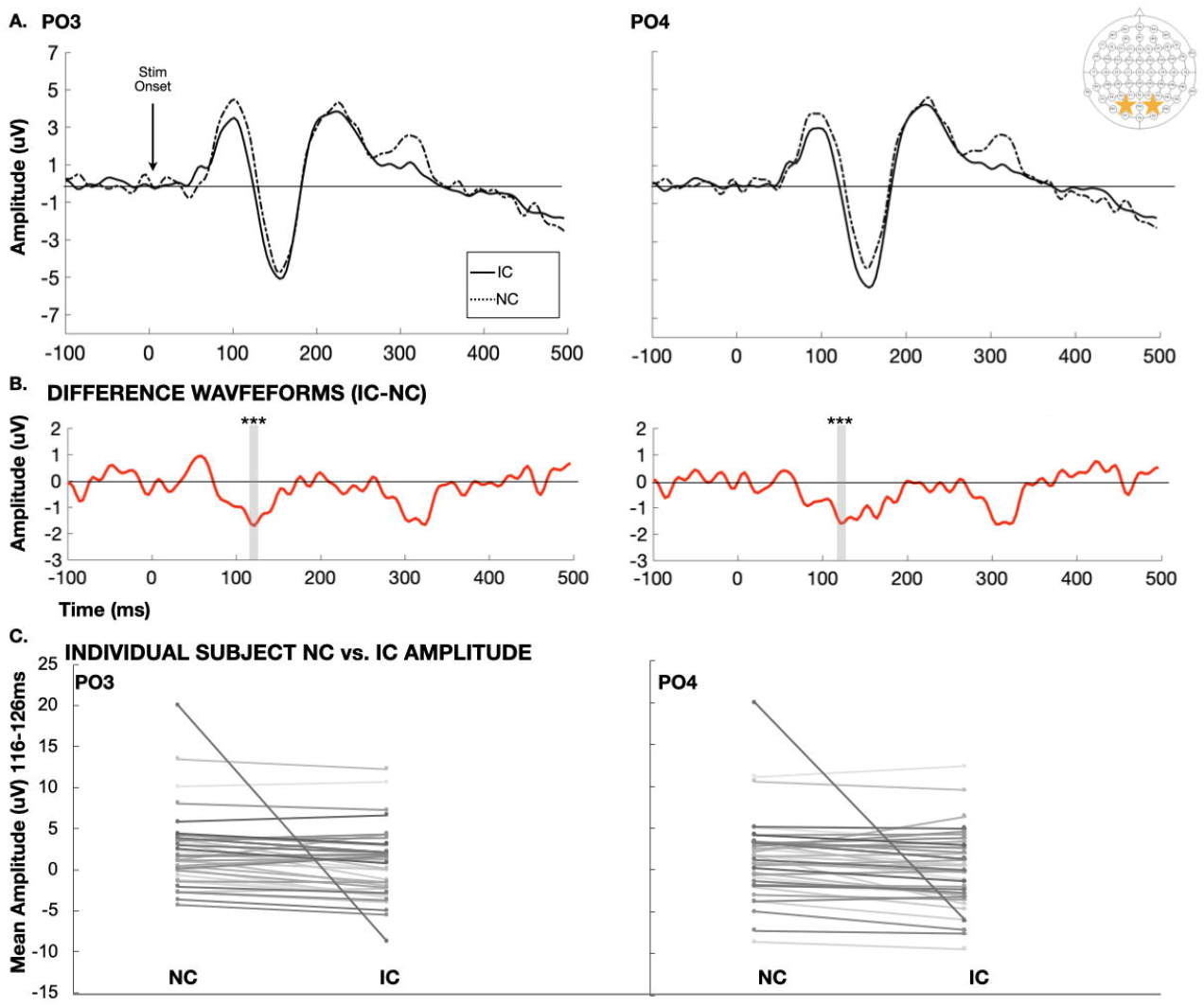
**A)** Grand-average VEP waveforms across all participants (irrespective of behavioral performance) elicited by the IC (solid) and NC (dotted line) stimuli over the left and right lateral occipital regions (PO3 and PO4 in the 10-20 system convention). Electrode positions marked by stars in the scalp schematic shown on the top right corner. B) Difference waveforms representing the evoked response to IC minus NC stimuli at electrodes PO3 (left) and PO4 (right). Time window for the whole group analysis (116-126ms) marked by the shaded region. *** p<.001 C) Individual subject mean amplitudes evoked by NC and IC stimuli in the IC-effect time window (116-126ms) at electrodes PO3 (left) and PO4 (right).

#### Relationship between Individual IC-effect and Behavioral Performance

Figure 4 displays for each level of performance (below average, low average, high average, and superior) grand-average VEP waveforms elicited by the IC and NC stimuli (Figure 4A), as well as the corresponding difference waves (Figure 4B), over the left (PO3) and right (PO4) occipital region. Topographic maps demonstrate the bilateral occipital negative foci, which are of clearly higher amplitude among participant groups with higher levels of performance as compared to those with lower levels of performance (Figure 5). To test the hypothesis that the robustness of IC processing would be associated with the level of behavioral resistance to backward masking, we then defined a time window of 10ms centered around each individual participant’s IC-effect negative peak latency between 100ms-200ms to obtain a mean amplitude for the IC-effect difference waveform for each individual. A one-way ANOVA using these mean amplitude measures as the dependent measure revealed significant differences between groups in their IC-effect (F(3,36)=3.498, p=0.025, η_p_^2^=0.226) and post-hoc analyses confirmed significantly greater IC-effect amplitude in superior as compared to below average performance groups, as well as a non-significant trend in the same direction when comparing superior performers to each of the high and low average performance groups (Table 1). One-way ANOVA comparing group classification with negative peak latency (between 100-200ms) did not reveal group differences (F(3,36)=0.475, p=0.702, η_p_^2^=0.038), suggesting that timing of the IC-effect was not a key factor in the current dataset.

**Figure 4.**
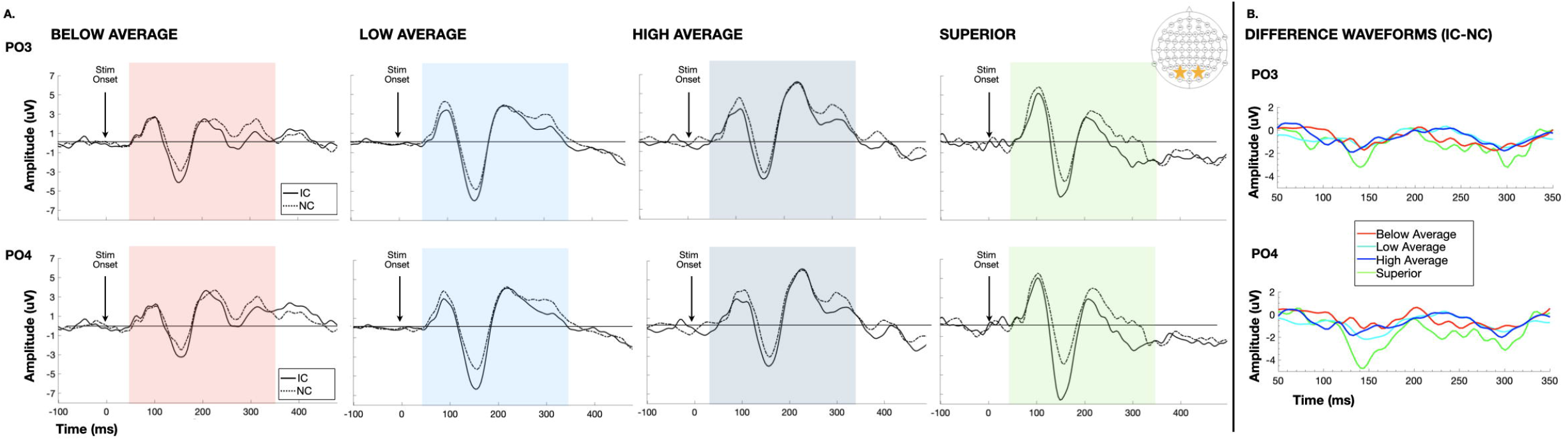
**A)** Grand-average VEP waveforms elicited by the IC (solid) and NC (dotted line) stimuli over the PO3 (top panel) and PO4 (bottom panel) electrode locations for participants grouped by performance on the behavioral backward masking paradigm (below average-red, low average-light blue, high average-dark blue, superior-green). Electrode positions marked by stars in the scalp schematic shown on the top right corner. Colored regions of shading mark the time period for which differences waveforms are displayed in panel B. **B)** Difference waveforms representing the evoked response to IC minus NC stimuli at electrodes PO3 (top panel) and PO4 (bottom panel) for below average (red), low average (light blue), high average (dark blue), and superior (green) performers.

**Figure 5.**
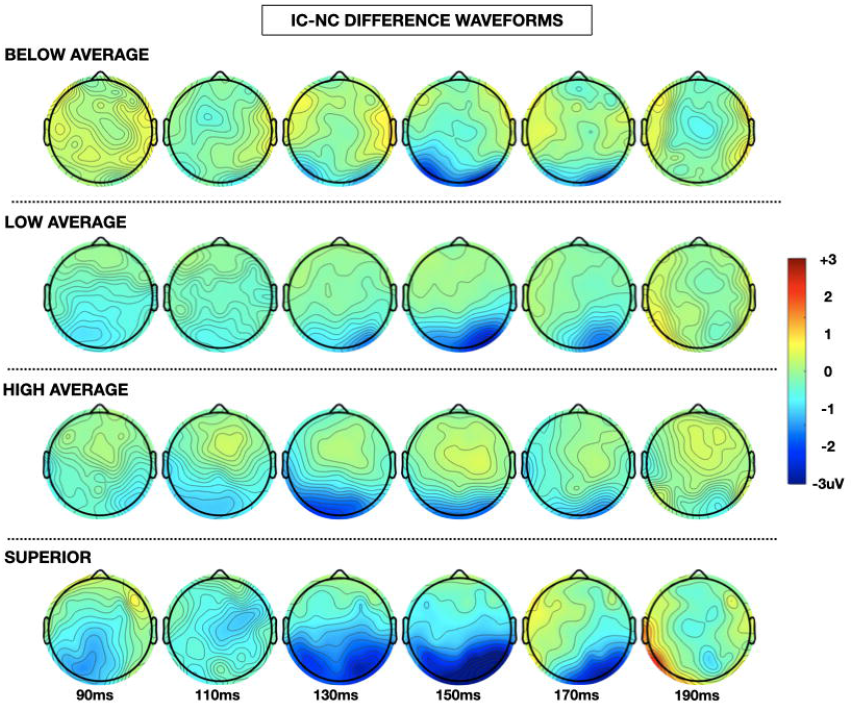
Topographic representation of the difference in instantaneous amplitude of evoked response between IC and NC stimuli at 20ms intervals between 90-190ms post-stimulus onset for participants grouped by performance (below average, low average, high average, superior)

**TABLE 1.**
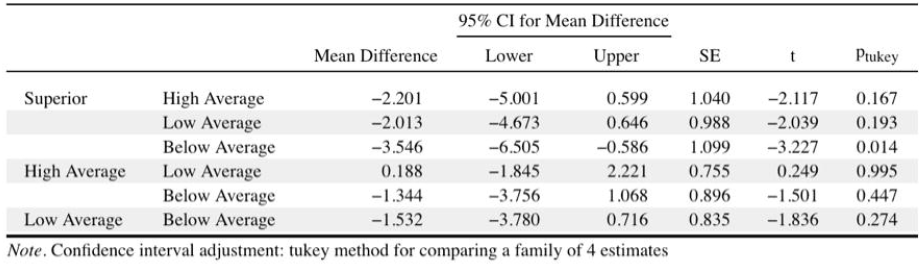

|  |  |  | 95% CI for Mean Difference |  |  |  |  |
| --- | --- | --- | --- | --- | --- | --- | --- |
|  |  | Mean Difference | Lower | Upper | SE | t | Ptukey |
| Superior | High Average | −2.201 | −5.001 | 0.599 | 1.040 | −2.117 | 0.167 |
|  | Low Average | −2.013 | −4.673 | 0.646 | 0.988 | −2.039 | 0.193 |
|  | Below Average | −3.546 | −6.505 | −0.586 | 1.099 | −3.227 | 0.014 |
| High Average | Low Average | 0.188 | −1.845 | 2.221 | 0.755 | 0.249 | 0.995 |
|  | Below Average | −1.344 | −3.756 | 1.068 | 0.896 | −1.501 | 0.447 |
| Low Average | Below Average | −1.532 | −3.780 | 0.716 | 0.835 | −1.836 | 0.274 |

*Note.* Confidence interval adjustment: tukey method for comparing a family of 4 estimates

Finally, the relationship between participant performance as a continuous variable and IC-effect mean amplitude revealed significant correlation between an individual’s accuracy in the backward masking task at the 4 frame (r(38) = −0.433, p=0.003, one-tailed), 7 frame (r(38) = −0.410, p=0.004, one-tailed), and 10 frame (r(38) = −0.324, p=0.021, one-tailed) target-to-mask delay conditions. There was also a non-significant trend toward association between greater IC-effect mean amplitude and performance at the 1 frame delay (r(38) = −0.222, p=0.085, one-tailed) and no mask conditions (r(38) = −0.227, p=0.079, one-tailed). Together, performance across all five target-to-mask delay conditions accounted for approximately 27% of the variance in IC-effect mean amplitude (F(5,34) =2.494, p=0.050, R^2^=0.268, see Figure 6).

**Figure 6.**
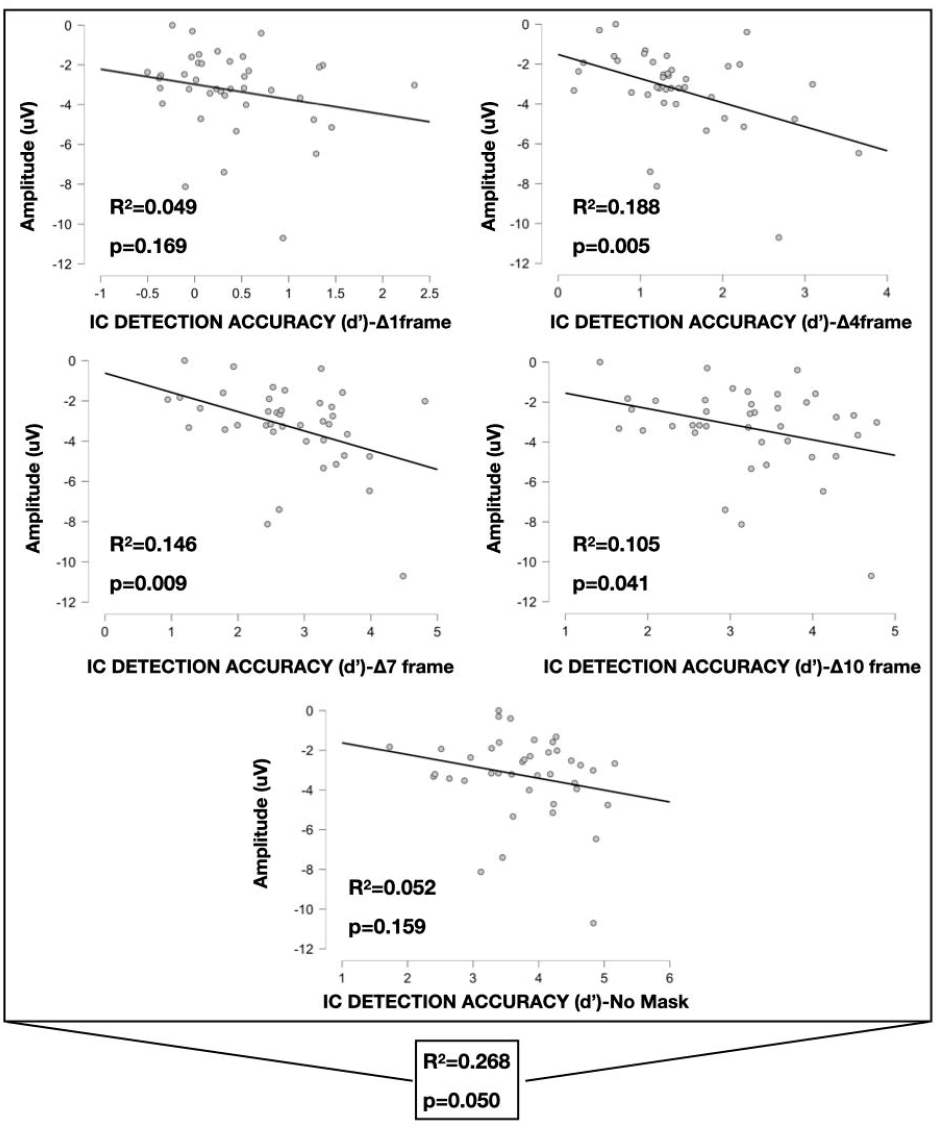
Scatter plot depicting the relationship between each individual’s maximal difference in IC-NC mean amplitude (uV) during the general *IC-effect time* window (100ms-200ms) and performance on the behavioral backward masking paradigm at each of the five target-to-mask delay conditions (1, 4, 7, 10 frames and no mask). Linear regression suggests performance across all five stimulus-mask delay conditions accounts for approximately 27% of the variance in IC-effect mean amplitude (F(5,34) =2.494, p=0.050, R^2^=0.268).

## DISCUSSION

Consistent with a large body of prior work (2–4, 6, 38), presentation of IC stimuli generated a clear negative-going modulation of the VEP in the time-period around the N1 component (~100-200ms post-stimulus onset) relative to the VEP response to NC stimuli. This “IC-effect” showed variability across individuals in terms of its amplitude during an electrophysiological recording session where participants were engaged in a central fixation task and simply ignored the task-irrelevant IC/NC inducer stimuli. In a separate experimental session, the same participants were asked to detect the presence or absence of IC stimuli under variable backward masking conditions, where the time-gap between the inducers and subsequent mask was varied parametrically. Here again, a large degree of inter-individual variability was observed in terms of task performance. This allowed us to stratify our cohort into four groups ranging from poor performers to superior performers; that is those who were very susceptible to the effects of the masking input versus those who were highly resistant to its effects, and the gradations in between. The data were consistent with our main hypothesis that the strength of the IC-effect would be a predictor of performance in the separately conducted backward masking psychophysical task, with a significant relationship between the VEP measures and task performance observed using correlational analysis. The data support the thesis that the strength of feedback processing is an indicator of the speed and robustness by which visual perception occurs in humans, at least for this class of stimuli.

It is important to point out that the current work does not argue that feedback processes occur only after some discrete and protracted delay. Such a simplistic hierarchical model is simply not plausible based on anatomy, connectivity patterns and the temporal processing dynamics of various cell types in the system. Indeed, it is clear from intracranial work in non-human primates that feedback processes begin extremely early in processing (39), and that recurrent processing across the entire visual system occurs within a matter of 10-25ms (40, 41). Therefore, variation in timing and efficiency of communication across different nodes within these networks may account for individual variability in susceptibility to disruption of contour integration by subsequent presentation of a masking stimulus.

To our knowledge, this is the first study in humans to demonstrate a relationship between the magnitude of this IC-related VEP modulation and inter-individual variability in susceptibility to disruption of contour visual discrimination by subsequent presentation of a random noise mask. Individuals with superior performance on the behavioral backward masking paradigm assessing IC vs. NC discrimination demonstrated an IC-effect that was greater in magnitude in the separate electrophysiologic experiment, indicating that individuals with more robust IC-related VEP modulation were more resistant to backward masking interference. This finding is consistent with non-human primate studies demonstrating that specific modulation of the underlying electrophysiologic processing in primary visual cortex directly impacted behavioral performance on a figure-ground discrimination task (9, 27, 29, 30). Although one might also predict that resistance to masking at shorter target-to-mask intervals should require more rapid processing of the IC/NC stimuli, performance in the current study was not significantly related to the latency of the IC-effect. We would caution though that it would be premature to rule out a timing component to these processes on the basis of the current dataset, since a study in 40 participants may not be sufficiently powered to uncover small effects, and a quick inspection of the topographic maps in Figure 5 suggests that the timing of the IC-effect may be later in poorer performing individuals. Nonetheless, the main analysis here suggested that timing was not a major factor and that among the participants in this study, the most consistent factor supporting IC discrimination despite subsequent visual interference is the strength of the IC-related VEP modulation. Thus, individuals with improved performance are likely able to recruit sufficient neural units engaged in visual spatial processing prior to mask onset, conferring greater resistance to the mask.

These studies form a foundation for future investigation of the real-world functional significance of these feedback-related differences in both typical development and psychopathology. IC-related processing has been described in neurotypical children as young as 5 years of age and can be tracked developmentally across childhood (42). Delays in the onset of these processes have been observed in young adults with schizophrenia (13), although the robustness of these modulations does not appear to be a factor (2). Understanding the relationship between feedback timing and overall efficiency of visual processing across multiple types of visual-spatial tasks, as well as other multisensory processing or cognitive tasks may therefore have implications for early detection of individuals at risk for psychosis and/or monitoring of targeted intervention. However, additional investigation into the strength and consistency of these electrophysiologic-behavioral relationships across age and different clinical populations would be necessary to validate any potential utility as biomarkers of typical versus atypical development, as it is possible that the variability in IC-related feedback processing within a typical population as demonstrated in this study may limit the ability to distinguish psychopathology.

### Study Limitations

A series of post-hoc analyses were conducted to elucidate the relationship between robustness of the early illusory-contour-related VEP modulation and resistance to the effects of rapid backward masking by grouping into performance levels. As a result, the findings should be interpreted as exploratory and reveal interesting potential effects that generate testable hypotheses for validation in future large independent samples of typical and clinical populations. Additionally, there were larger numbers of participants with low and high average performance than with significantly below average or superior performance. While this is expected given that many continuous performance measures conform to a normal distribution, ability to conduct adequately powered group-level statistical analyses accounting for individuals in the tails of the distribution is limited by these sample sizes. Furthermore, within group variability may make it difficult to directly predict an individual’s performance solely on the basis of his/her electrophysiologic signature. In this study, just over 1/4 of the variability in degree of IC-related VEP modulation was accounted for by task performance. It is possible that controlling for other factors would help to further clarify some of the residual ambiguity. For example, there is currently a debate on the influence of attention on illusory contour perception whereby the response to illusory contours has generally been considered to be pre-attentive (43, 44); however, more recent investigations have suggested that attention may indeed play a role (45). While we attempted to limit attention-related effects through the use of eye tracking for fixation, we did not directly assess for inattention-related symptoms in our study population. Finally, VEPs were specifically assessed separately rather than during the backward masking behavioral paradigm in order to differentiate VEP modulation related specifically to IC discrimination from effects of the mask or cognitive demands of the task. A future potential question of interest is whether the degree of mask-related modulation of the IC-effect also relates to task performance, as mask-related modulation of the VEP has been demonstrated previously (22, 46).

### Conclusions

We demonstrated a relationship between the strength of visual feedback processing during illusory contour processing and inter-individual variability in task performance under visual backward masking conditions, whereby individuals with more robust IC-related modulation of the VEP are more resistant to the effects of rapid backward masking during illusory contour discrimination. This advances our understanding of the contribution of feedback processing to individual differences in sensory-perceptual function and lays the groundwork for future studies of the implications of inter-individual variability in these functions.

## Acknowledgements

We thank Mr. Eric Nicholas for his invaluable contributions to the experimental setup. Our thanks as well to Dr. Holger Sperdin (@HolgeregloH) and Dr. Bob Sekuler (@robsek) for providing some historical background on backward masking through the Twittersphere.

## Supplementary Files

Upon acceptance for publication, the authors will coordinate with the editorial office to ensure that full and appropriately de-identified datasets are uploaded to a publicly available repository and that the appropriate link is made to the article file.

## Ethics and Consent

The Research Subjects Review Board of the University of Rochester approved all the experimental procedures. Each participant provided written informed consent in accordance with the tenets laid out in the Declaration of Helsinki.

## Funding Information

This work was supported by the Ernest J. Del Monte Institute for Neuroscience Pilot Program with funding from the Kilian J. and Caroline F. Schmitt Foundation (to EGF and JJF). Participant recruitment and phenotyping were conducted through the UR-IDDRC Human Clinical Phenotyping and Recruitment Core, and electrophysiological recordings and eye-tracking were supported by the UR-IDDRC Translational Neuroimaging and Neurophysiology Core, which are supported by a center grant from the Eunice Kennedy Shriver National Institute of Child Health and Human Development (P50 HD103536 – to JJF). Dr. Knight was supported by the University of Rochester Medical Center Department of Pediatrics Chair Fellow Award and with a Kyle Family Fellowship. Dr. Evans was supported by a post-doctoral training fellowship through the Center for Visual Sciences at the University of Rochester (NIH T32 EY007125).

## Authors’ Contributions

JJF and EGF conceived of and designed the study. EJM and ZC implemented the paradigm, recruited and phenotyped the participants, conducted the original pilot work, and collected the data. EJM and ZC also conducted initial analyses. EJK conducted the main analyses, created the figures and conducted the statistical analyses, all in close consultation with JJF, EGF, and SM. JJF wrote the main text with substantial input from EJK, SM and EGF, and received multiple rounds of editorial input from all of the authors.

## Competing Interests

All authors affirm that they have no financial interests or other potential conflicts of interest to report.

## Abbreviations List

VEP: Visual Evoked Potential
LOC: Lateral Occipital Complex
IC: Illusory Contour
NC: No Contour
EEG: Electroencephalographic
ANOVA: Analysis of Variance

## Notes

### Competing Interest Statement

The authors have declared no competing interest.

