## Supplemental Materials for "The strength of feedback processing is associated with resistance to visual backward masking during illusory contour processing in adult humans"

**Table S1.** Numbers of trials included in the analysis across condition (IC/NC) in each of the four performance-related groups.

|  | **Below Average** | **Low Average** | **High Average** | **Superior** | **All Participants** |
| --- | --- | --- | --- | --- | --- |
| Avg. Accepted  NC trials ± SD  (Total trials≈  210) | 136±69 | 165±28 | 152±36 | 142±46 | 153±43 |
| Avg. Accepted IC trials ± SD  (Total trials≈  210) | 135±71 | 165±28 | 149±41 | 135±56 | 151±46 |
| Median Channels Interpolated (range) | 6 (1-10) | 3 (1-10) | 2 (1-7) | 1 (1-6) | 3 (1-10) |

**Figure S1.** Box plot depicting the age distribution in each of the four performance-related groups. All data points included in the analyses ^o^denotes group outliers (=3rd quartile + 1.5*interquartile range); *denotes extreme group outliers (=3rd quartile + 3*interquartile range).

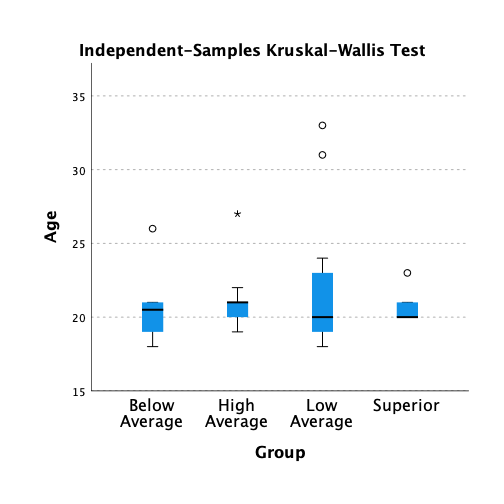
